# Compensatory epistasis maintains ACE2 affinity in SARS-CoV-2 Omicron BA.1

**DOI:** 10.1101/2022.06.17.496635

**Authors:** Alief Moulana, Thomas Dupic, Angela M. Phillips, Jeffrey Chang, Serafina Nieves, Anne A. Roffler, Allison J. Greaney, Tyler N. Starr, Jesse D. Bloom, Michael M. Desai

## Abstract

The Omicron BA.1 variant emerged in late 2021 and quickly spread across the world. Compared to the ancestral Wuhan Hu-1 strain and other pre-Omicron SARS-CoV-2 variants, BA.1 has many mutations, a number of which are known to enable antibody escape ^1–3^. Many of these antibody-escape mutations individually decrease the spike receptor-binding domain (RBD) affinity for ACE2 in the background of early SARS-CoV-2 variants ^4^, but BA.1 still binds ACE2 with high affinity ^5,6^. The fitness and evolution of the BA.1 lineage is therefore driven by the combined effects of numerous mutations. Here, we systematically map the epistatic interactions between the 15 mutations in the RBD of BA.1 relative to the Wuhan Hu-1 strain. Specifically, we measure the ACE2 affinity of all possible combinations of these 15 mutations (2 ^15^ = 32,768 genotypes), spanning all possible evolutionary intermediates from the ancestral Wuhan Hu-1 strain to BA.1. We find that immune escape mutations in BA.1 individually reduce ACE2 affinity but are compensated by epistatic interactions with other affinity-enhancing mutations, including Q498R and N501Y. Thus, the ability of BA.1 to evade immunity while maintaining ACE2 affinity is contingent on acquiring multiple interacting mutations. Our results implicate compensatory epistasis as a key factor driving substantial evolutionary change for SARS-CoV-2 and are consistent with Omicron BA.1 arising from a chronic infection.

---

The Omicron BA.1 variant of SARS-CoV-2 emerged in November 2021 and spread rapidly throughout the world, driven in part by its ability to escape existing immunity in vaccinated and previously infected individuals ^7,8^. Strikingly, Omicron did not emerge as a descendant of the then-widespread Delta lineage. Instead, it appeared as a highly diverged strain after accumulating dozens of mutations within a lineage that was not widely circulating at the time, including 15 mutations within the spike protein receptor-binding domain (RBD) ^7^.

Recent work has shown that a number of these 15 RBD mutations (some of which are seen in other variants) disrupt binding of specific monoclonal antibodies ^1,3,9–11^, potentially contributing to immune escape. However, most of these mutations have also been shown to reduce binding affinity to human ACE2 when they arise within the Wuhan Hu-1, Delta, or several other SARS-CoV-2 lineages ^4,12^, potentially impairing viral entry into host cells. In contrast, the Omicron RBD tolerates these escape mutations while retaining strong affinity to ACE2, suggesting that other mutations in this lineage may help maintain viral entry.

Earlier work has systematically analyzed mutational effects on antibody binding and ACE2 affinity, for example by using deep mutational scanning (DMS) ^12,13^. However, these approaches focus on the effects of single mutations on specific genetic backgrounds. They are therefore useful for understanding the first steps of evolution from existing variants but cannot explain how multiple mutations interact over longer evolutionary trajectories. Thus, it remains unclear how combinations of mutations, such as those observed in Omicron, interact to both evade immunity and retain strong affinity to ACE2.

To address this question, we used a combinatorial assembly approach to construct a plasmid library containing all possible combinations of the 15 mutations in the Omicron BA.1 RBD (a total of 2 ^15^ = 32,768 variants). This library includes all possible evolutionary intermediates between the Wuhan Hu-1 and Omicron BA.1 RBD. We transformed this plasmid library into a standard yeast display strain, creating a yeast library in which each cell displays a single monomeric RBD variant corresponding to the plasmid in that cell. We then used Tite-Seq, a high-throughput flow cytometry and sequencing-based method ^14,15^ (see Methods; Extended Data Figure 1A), to measure the binding affinities, *K*_D,app_, of all 32,768 RBD variants to human ACE2 in parallel. Consistent with earlier work by ourselves ^15^ and others ^12,14,16^, we find that these Tite-Seq measurements are highly reproducible (SEM of 0.2 log *K*_D,app_ between triplicate measurements) and consistent with independent low-throughput measurements (see Methods; Extended Data Figure 1B-F). In addition, we find minimal variation in RBD expression levels and are thus able to infer *K*_D,app_ for the entire combinatorial library (see Methods; Extended Data Figure 2).

We find that all 32,768 RBD intermediates between Wuhan Hu-1 and Omicron BA.1 have detectable affinity to ACE2, with *K*_D,app_ ranging between 0.1 μM and 0.1 nM (Figure 1A and Extended Data Figure 1; see https://desai-lab.github.io/wuhan_to_omicron/ for an interactive data browser). Consistent with previous studies ^5^, the BA.1 RBD exhibits a slight (3-fold) improvement in binding affinity compared to Wuhan Hu-1. However, most (~ 60%) of the intermediate RBD sequences actually show a weaker binding affinity to ACE2 than the ancestral Wuhan Hu-1 RBD. This is because the vast majority of BA.1 mutations have a neutral or deleterious effect on ACE2 affinity on most genetic backgrounds (Figure 1B). This is particularly true for K417N, G446S, Q493R, G496S, and Y505H, four of which are known to be involved in escape from various classes of monoclonal antibodies ^17–19^.

**Figure 1.**
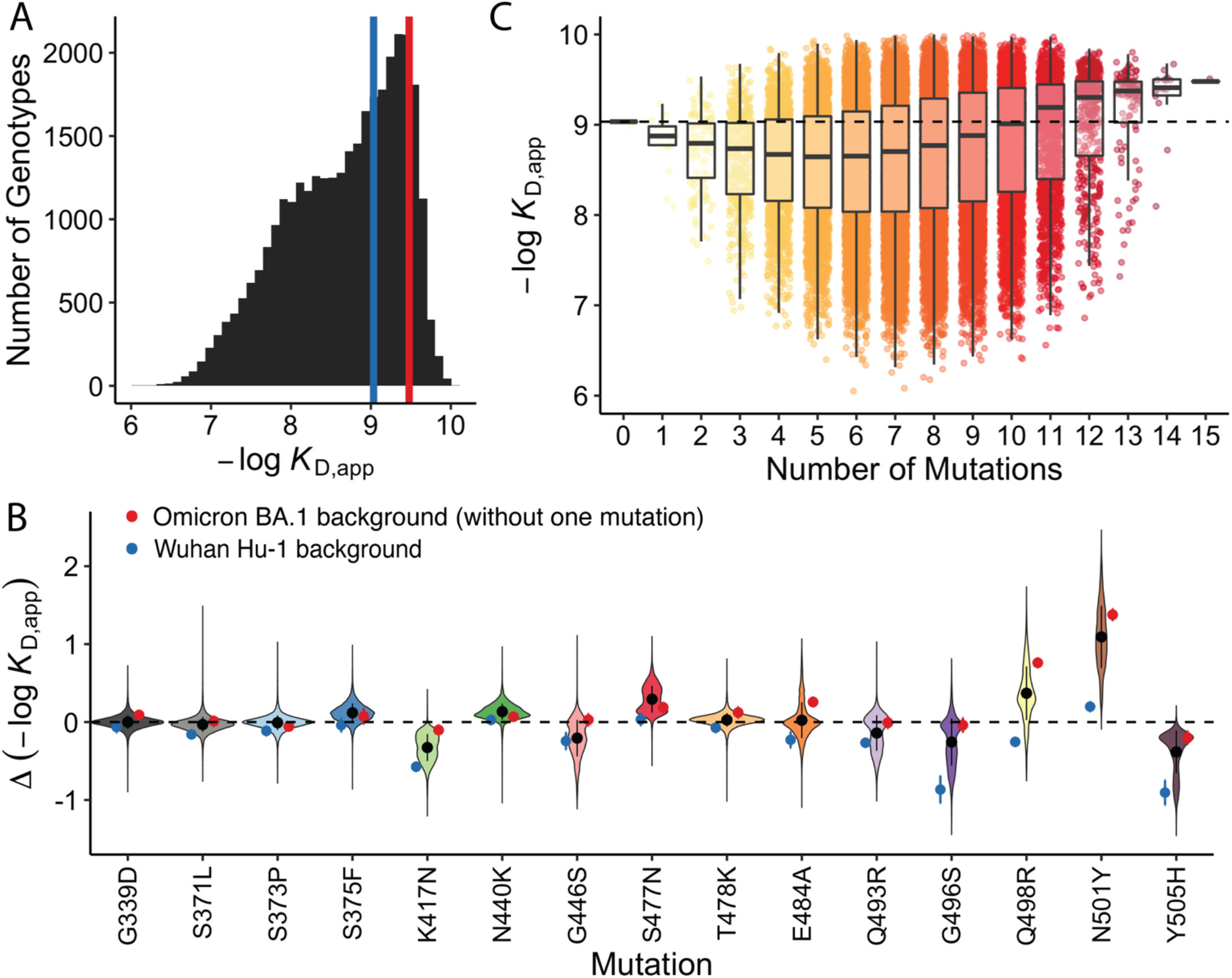
Binding affinity landscape. (**A**) Distribution of binding affinities to ACE2 across all N=32,768 RBD genotypes tested. Binding affinities are shown as -log*K*_D,app_; vertical blue and red lines indicate the - log*K*_D,app_ for Wuhan Hu-1 and Omicron BA.1, respectively. (**B**) Distributions of the effect of each mutation on ACE2 affinity (defined as the change in -log*K*_D,app_ resulting from mutation) across all possible genetic backgrounds at the other 14 loci. Black line segments indicate 25th and 75th percentiles of the effect distributions and points represent distribution means. Blue and red points specify effects on Wuhan Hu-1 and Omicron BA.1 backgrounds, respectively. (**C**) Distribution of binding affinities grouped by number of Omicron BA.1 mutations. Binding affinity of the Wuhan Hu-1 variant is indicated by horizontal dashed line.

Although many BA.1 mutations reduce ACE2 affinity on average, the interactions between these mutations result in improvement in ACE2 affinity for BA.1 relative to the ancestral Wuhan Hu-1 strain. That is, mutations tend to be more deleterious for ACE2 affinity if few other mutations are present but tend to become neutral or even beneficial in the presence of multiple other mutations (Figure 1C; Extended Data Figure 3). Consistent with this, we find that although most of the 15 RBD mutations reduce ACE2 affinity in the Wuhan Hu-1 background (and in many cases across most other backgrounds as well), they all become less deleterious or even beneficial in the Omicron background (Figure 1B). This pattern explains why the BA.1 RBD has a stronger affinity for ACE2 despite containing so many mutations that individually reduce ACE2 affinity: their deleterious effects are mitigated by compensatory epistatic interactions with other mutations.

To systematically analyze mutational effects and interactions, we fit a standard biochemical model of epistasis ^20^ to our data. This decomposes our measured -log(*K*_D,app_) (which is expected to be proportional to the free energy of binding, ΔG) ^21,22^ into a sum of linear effects from single mutations, pairwise epistasis, and higher-order epistatic interactions among larger sets of mutations (truncated at fifth order; Extended Data Figure 4, see Methods). This model yields coefficients that are comparable to alternative models of statistical (Extended Data Figure 5) and global ^23^ (Extended Data Figure 6) epistasis. Generally, we find that the linear effects of individual mutations (Figure 2A) correlate with the ACE2 contact surface area of the corresponding residue (Figure 2B,C), and neighboring residues are more likely to have strong pairwise interactions (Figure 2E), as we might expect from previous work ^15,24^.

**Figure 2.**
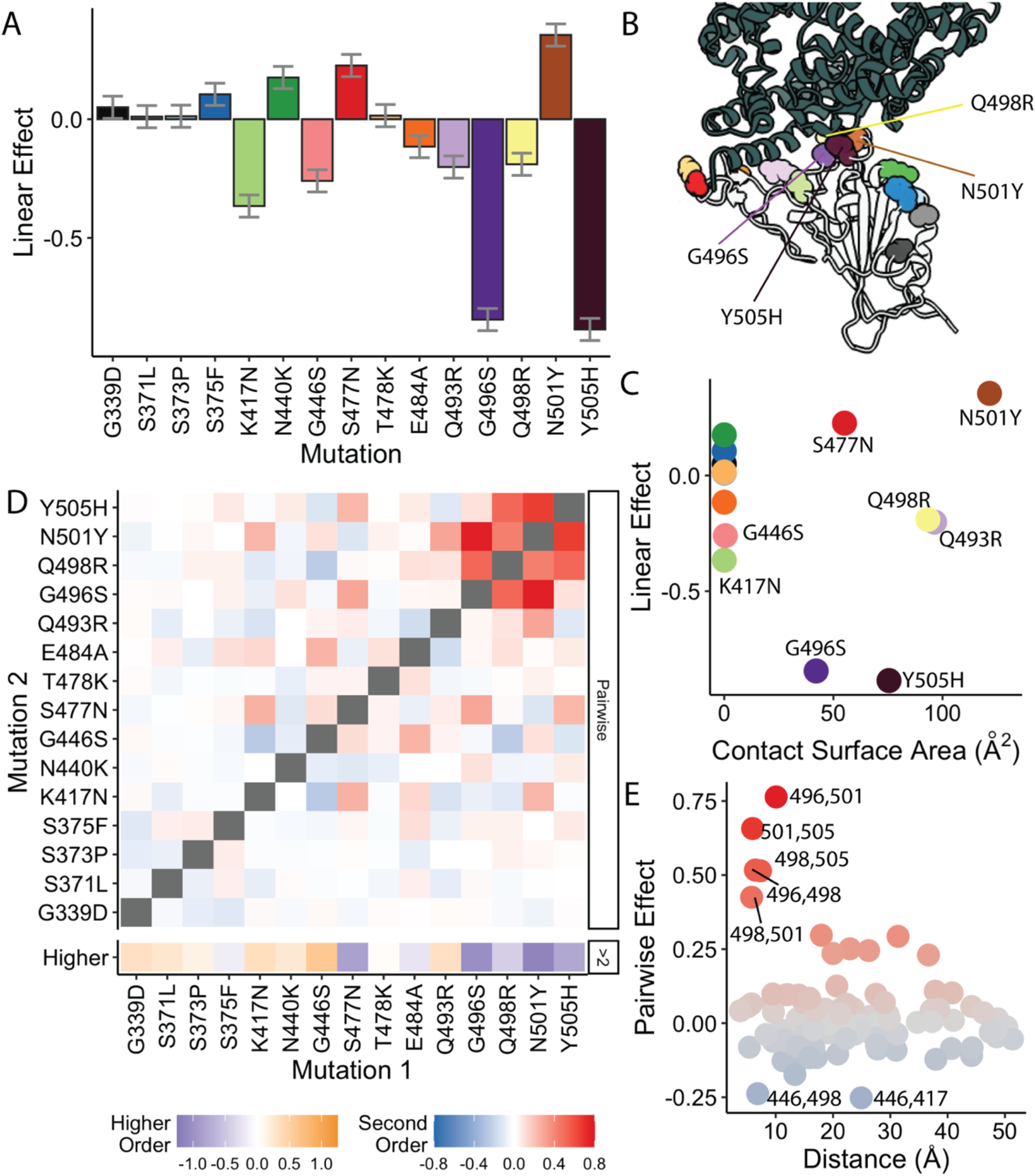
Linear and epistatic effects of mutations. (**A**) First-order effects in best-fitting epistatic interaction model (up to fifth order). Error bars represent standard errors from the model fit. (**B**) Co-crystal structure of Omicron BA.1 RBD and ACE2 receptor (PDB ID 7WPB). Mutated residues shown as spheres colored as in (**A**). (**C**) First-order effects for each mutation plotted against contact surface area between corresponding BA.1 RBD residue and ACE2. Mutations colored as in (**A**). (**D**) Second-order epistatic interaction coefficients and higher order interaction coefficients. For each mutation, higher order interaction coefficient (shown at bottom of heat map plot) is calculated by summing over all third- and fourth-order interaction coefficients involving the mutation. (**E**) Pairwise interaction coefficients plotted against the distances between the respective alpha-carbons. Mutations are colored by pairwise coefficient as in (**D**).

Our inferred pairwise and higher-order coefficients reveal that strong compensatory interactions offset the effects of affinity-reducing mutations (Figure 2D). The magnitude of these interactions is comparable to that of the linear effects, and this epistasis is overwhelmingly positive, as excluding epistatic terms leads to a consistent underestimate of the predicted affinity (Extended Data Figure 7). This strong positive epistasis means that mutations which reduce ACE2 affinity become less deleterious in backgrounds containing other compensatory mutations. For example, the negative linear effect of Q498R is fully compensated by its interaction with nearby mutation N501Y; this pairwise interaction has been highlighted in earlier work ^4,6,25^. We identify numerous additional interacting mutations, including even stronger positive interactions (along with third and fourth-order effects) between Q498R, G496S, N501Y, and Y505H (Figure 2D).

This high-order compensatory epistasis eliminates the strongly deleterious effects of mutations involved in antibody escape on ACE2 affinity. Specifically, earlier work has shown that five BA.1 mutations (K417N, G446S, E484A, Q493R, and G496S) have a particularly strong effect in promoting antibody escape ^1,18,19^. These mutations all individually reduce affinity to ACE2 both on average and in the Wuhan Hu-1 background (except E484A; Figure 1B, 2A, 3A), and the combination of all five is strongly deleterious (Figure 3A,B). However, strong high-order epistasis with Q498R and N501Y mitigates this: either N501Y or Q498R alone reduces the cost of the five escape mutations, and the combination of both almost fully compensates for these deleterious effects (Figure 3B). While these escape mutations do also benefit from interactions with other mutations (Extended Data Figure 8), N501Y and Q498R account for the majority of the compensatory effect. We note that strong compensatory interactions also mitigate the deleterious effect of Y505H (Figure 3C). This mutation has not previously been shown to be strongly involved in antibody escape, but the pattern of compensation we observe suggests that it may be functionally relevant in some way.

**Figure 3.**
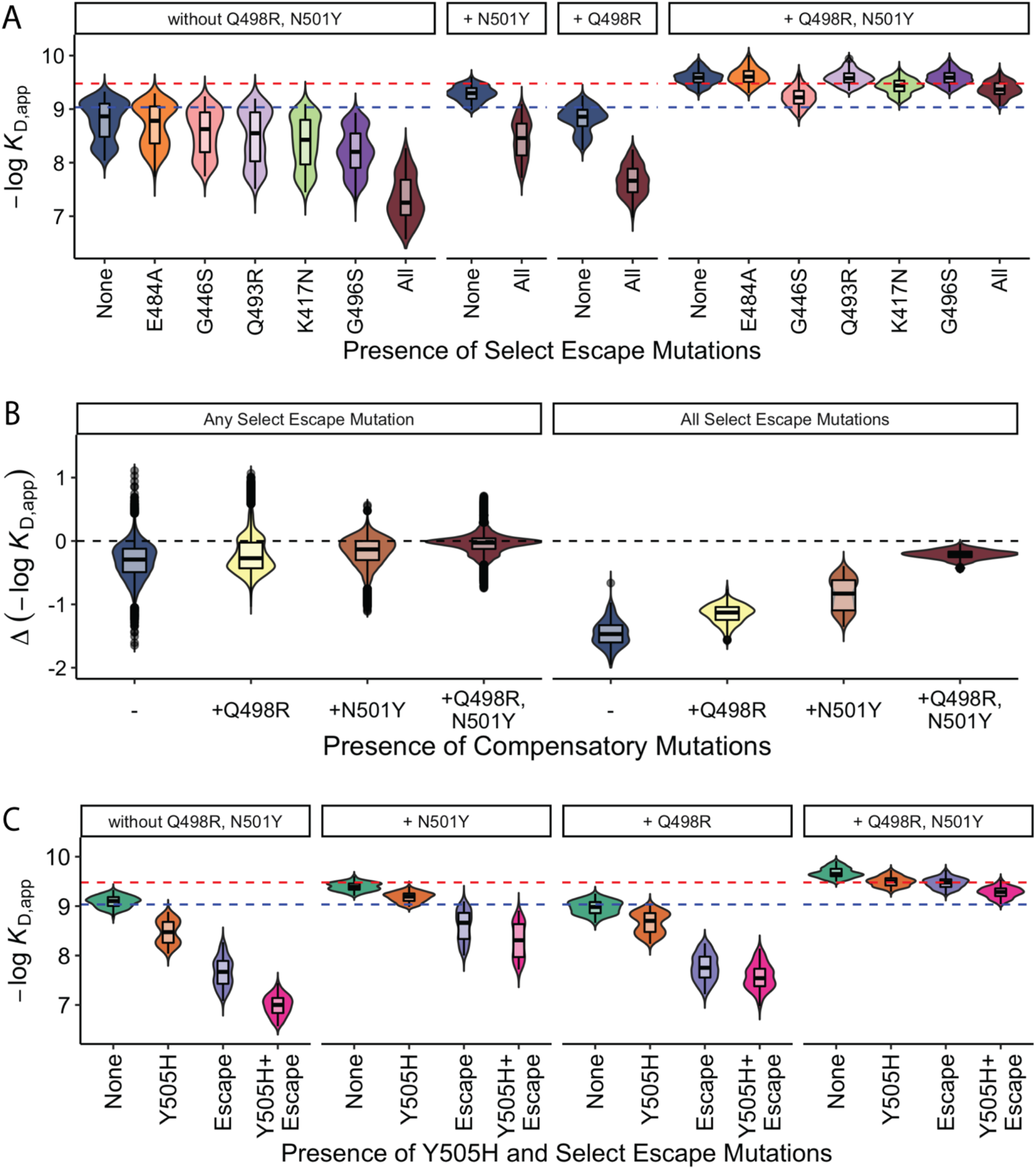
Epistasis compensates for reductions in ACE2 affinity. (**A**) ACE2 binding affinities for variants containing mutations that have a strong effect on antibody escape: K417N, G446S, E484A, Q493R, and G496S grouped by the presence of compensatory mutations (Q498R and N501Y). Dashed blue (resp. red) line indicates Wuhan Hu-1 (resp. Omicron BA.1) ACE2 binding affinity. (**B**) The changes in ACE2 binding affinities for variants containing any one (or all) of select escape mutations grouped by the presence of compensatory mutations (Q498R and N501Y). Dashed line indicates no affinity change. (**C**) ACE2 binding affinities for variants containing Y505H and antibody escape mutations presented as in (**A**).

The extensive epistasis we observe means that the individual effects of each of these 15 mutations, as well as the pairwise interactions between them, are likely different in other viral lineages. However, earlier work has shown that the antibody escape mutations described above (K417N, G446S, E484A, Q493R, and G496S) similarly reduce ACE2 affinity in several other variants (including Alpha, Beta, Eta, and Delta) ^25^. Consistent with this result, we find that these mutations, along with others that we find have a negative linear effect on ACE2 affinity, rarely occur across the SARS-CoV-2 phylogeny (Figure 4A). This suggests that maintaining affinity to human ACE2 is likely an important aspect of viral fitness, so these mutations are typically selected against. Similarly, we find that mutations with negative effects on ACE2 affinity that are compensated by epistatic interactions with N501Y tend to be enriched across the SARS-CoV-2 phylogeny in strains that also have N501Y, relative to strains that do not (Figure 4B; other pairwise interactions co-occur too rarely to test). This further suggests that at least some of the pairwise epistatic interactions we observe are also present in other backgrounds, and that viral evolution has favored compensation for reduction in ACE2 affinity.

**Figure 4.**
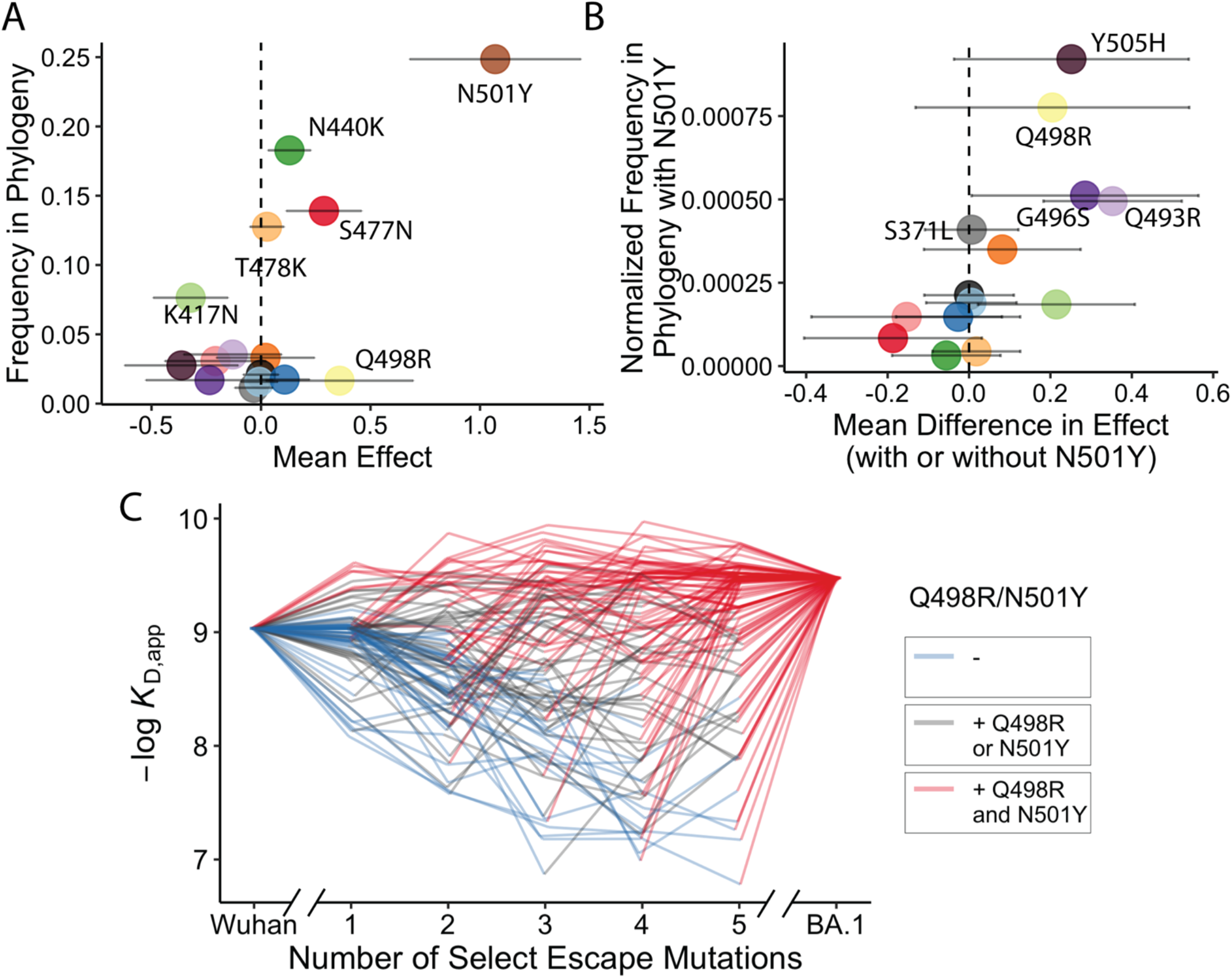
Trajectory of Omicron BA.1 evolution. (**A**) Frequency of occurrences for each mutation across SARS-CoV-2 sequences available on GISAID (see Methods) as a function of their average effect on ACE2 affinity in our data. Error bars indicate standard deviation of effect sizes. (**B**) Normalized frequency of mutations co-occurring with N501Y across SARS-CoV-2 sequences available on GISAID (calculated based on the frequency at which each mutation occurs on the same branch as N501Y, normalized by their overall frequency; see Methods) as a function of the difference in their effect on ACE2 affinity in the presence of N501Y. Error bars indicate standard deviation of effects. (**C**) ACE2 affinity trajectories for 100 randomly selected pathways (involving all 15 mutations), shown as a function of the number of mutations with strong effect on antibody escape (K417N, G446S, E484A, Q493R, and G496S) and the presence or absence of compensatory mutations Q498R and N501Y (shown with colors). Each trajectory represents a possible mutation order, starting at the Wuhan Hu-1 genotype and ending at Omicron BA.1.

Together, these results suggest that the evolution of antibody escape in BA.1 was possible without disrupting binding to ACE2 because of the compensatory interactions with numerous other mutations unique to this lineage. While signatures of these selection pressures and epistatic interactions are present across the viral phylogeny ^26^, and antibody escape variants could have been compensated by other combinations of mutations, it is only the BA.1 lineage which accumulated this particular combination of interacting compensatory mutations.

Our results also provide insight into why the immune escape phenotype observed in Omicron BA.1 did not arise as the result of mutations accumulating within the then-widely circulating Delta variant. Specifically, the combination of multiple mutations required for both immune escape and maintaining affinity to ACE2 (Figure 4C) is unlikely to have accumulated within the context of acute infections, which involve few mutations between transmission bottlenecks and presumably strong selection pressures on both functions ^27^. In contrast, in chronic infections (e.g. in an immunocompromised host) large population sizes and relaxed selection pressures may allow for the accumulation of the many mutations required to both maintain ACE2 affinity and evade neutralizing antibodies ^28,29^. Under such relaxed selection, the compensatory mutations may have preceded the immune escape mutations, minimizing their otherwise deleterious effects on ACE2 affinity. Alternatively, relaxed selection for binding ACE2 may have created a permissive environment for the immune escape mutations, followed by compensation that then allowed the variant to spread to other hosts. Phylogenetic analysis provides some support for the former possibility, as two immune escape mutations (G446S and G496S) occur late in BA.1 evolution (and are not shared with the BA.2 lineage; Extended Data Figure 9). In addition, a strong selection model based on ACE2 affinity prefers the three BA.1-specific mutations to appear late in the evolution, as observed in the phylogeny (Extended Data Figure 10). Irrespective of the exact order of mutations, the large viral population size and relaxed selection pressure of a chronic infection may have created conditions conducive to the fixation of the several mutations required for BA.1 to evade neutralizing antibodies while maintaining ACE2 affinity.

We emphasize that our work is confined to 15 mutations within a specific region of one protein, and hence neglects potential interactions with the many other mutations present in the Omicron BA.1 lineage. In addition, we focus on ACE2 affinity and antibody escape, which represent only two aspects of viral fitness. It is likely that other properties of BA.1 (e.g. spike protein expression and stability) also play key roles in viral evolution. We find some hints of this in our data. For example, we identify a significant synergistic interaction between S371L, S373P, and S375F that improves RBD expression in yeast, consistent with earlier work showing that this set of mutations is associated with stabilization of a more tightly packed down-conformation of the RBD ^30^ (Extended Data Figure 3). Beyond this, numerous other phenotypes are also likely to be relevant.

Despite these caveats, our results demonstrate that key events in viral evolution can depend on high-order patterns of epistasis. This may be especially important for complex adaptive events involving numerous mutations, such as immune escape and host-switching. Thus, to predict the future of viral evolution we must move beyond high-throughput screens of single mutations, and more comprehensively analyze combinatorial sequence space. A key challenge is the vastness of this sequence space, which makes exhaustive exploration intractable. However, generating specific combinatorial landscapes like those presented here may help reveal general patterns of epistasis that shape viral evolution in complex environments.

## METHODS

### Yeast display plasmid & strains

To generate clonal yeast strains for the Wuhan Hu-1 and Omicron BA.1 variants, we cloned the corresponding RBD gblock (IDT, https://github.com/desai-lab/compensatory_epistasis_omicron/tree/main/Supplementary_Files) into pETcon yeast surface-display vector (plasmid 2649; Addgene, Watertown, MA, #166782; https://github.com/desai-lab/compensatory_epistasis_omicron/tree/main/Supplementary_Files) via Gibson Assembly. The sequence of the gblock was codon-optimized for yeast (using the Twist Bioscience algorithm); we found that the codon optimization had a significant impact on display efficiency. Additionally, for the library construction (described below), we deleted two existing Bsa-I sites from the plasmid by site-directed mutagenesis (Agilent, Santa Clara, CA, #200521). In the clonal strain production, Gibson Assembly products were transformed into NEB 10-beta electrocompetent *E. coli* cells (NEB, Ipswich, MA, #C3020K), following the manufacturer protocol. After overnight incubation at 37°C, the cells were harvested, and the resulting plasmids were purified and Sanger sequenced. We transformed plasmids containing the correct sequences into the AWY101 yeast strain (kind gift from Dr. Eric Shusta) ^31^ as described by Gietz and Schiestl ^32^. Transformants were plated on SDCAA-agar (1.71 g/L YNB without amino acids and ammonium sulfate [Sigma-Aldrich #Y1251], 5 g/L ammonium sulfate [Sigma-Aldrich #A4418], 2% dextrose [VWR #90000–904], 5 g/L Bacto casamino acids [VWR #223050], 100 g/L ampicillin [VWR #V0339], 2% Difco Noble Agar [VWR #90000–774]) and incubated at 30°C for 48 hr. Several colonies were restreaked on SDCAA-agar and again incubated at 30°C for 48 hr. Clonal yeast strains were picked, inoculated, grown to saturation in liquid SDCAA (6.7 g/L YNB without amino acid VWR #90004-150), 5 g/L ammonium sulfate (Sigma-Aldrich #A4418), 2% dextrose (VWR #90000– 904), 5 g/L Bacto casamino acids (VWR #223050), 1.065 g/L MES buffer (Cayman Chemical, Ann Arbor, MI, #70310), 100 g/L ampicillin (VWR # V0339)) at 30°C, and mixed with 5% glycerol for storage at -80°C.

### Yeast display library production

We generated the RBD variant library with a Golden Gate combinatorial assembly strategy. First, we divided the RBD sequence into five fragments of about equal length, ranging from 90 to 131 bp and each containing between 1 and 4 mutations. We introduced BsaI sites and overhangs to both ends of each fragment sequence. These overhangs contained BsaI cut sites that would allow the five fragments to assemble uniquely in their proper order within the plasmid backbone. For each fragment with *n* mutations, we generated 2 ^*n*^ fragment versions by either producing the fragments via PCR (Fragments 1-4) or purchasing individual DNA duplexes (Fragment 5) from IDT. These permutations ensured the inclusion of all possible mutation combinations in the library. In Fragment 2, we also included a synonymous substitution on the K378 residue that corresponds to the K417N mutation. This substitution allows for the amplicon library to be sequenced on the Illumina Novaseq SP (2×250bp). For dsDNA production by PCR, we designed the fragments such that the mutations they contain are close to the 3’ or 5’ ends. This design enabled the primers to simultaneously include and introduce the mutations, BsaI sites, and unique over-hangs chosen during the PCR. We produced each version of each fragment individually (28 PCR reactions in total; see https://github.com/desai-lab/compensatory_epistasis_omicron/tree/main/Supplementary_Files for primer sequences) and pooled the products of each fragment in equimolar ratios. Additionally, we also pooled all 16 purchased DNA duplexes encoding the fifth fragment in equimolar ratios. We then created a final fragment mix by pooling the five fragment pools together. In the Golden Gate reaction, the versions of each fragment would be ligated together in random combinations, producing all of the sequences present at approximately equal frequencies.

In addition to the fragment mix, we prepared four versions of the plasmid backbone for the Golden Gate reaction. Each version contains a combination of the mutations N501Y and Y505H. Prior to the assembly, we introduced the counter-selection marker *ccdB*, in place of the fragment insert region, with flanking BsaI sites (https://github.com/desai-lab/compensatory_epistasis_omicron/tree/main/Supplementary_Files). We performed Golden Gate cloning using Golden Gate Assembly Mix (NEB, Ipswich, MA, #E1601L), following the manufacturer recommended protocol, with a 7:1 molar ratio of the fragment insert pool to plasmid backbone. We transformed the assembly products into NEB 10-beta electrocompetent *E. coli* cells in 6 × 25 μL cell aliquots. We then transferred each of the recovered cell culture to 100 mL of molten LB (1% tryptone, 0.5% yeast extract, 1% NaCl) containing 0.3% SeaPrep agarose (VWR, Radnor, PA #12001– 922) spread into a thin layer in a 1L baffled flask (about 1 cm deep). The mixture was placed at 4°C for three hours, after which it was incubated for 18 hr at 37°C. We observed a total of 3 million transformants across aliquots. To isolate the plasmid library, we mixed the flasks by shaking for 1 hr and pelleted the cells for standard plasmid maxiprep (Zymo Research, Irvine, CA, D4201), from which we obtained > 90 μg of purified plasmid.

We then transformed the purified plasmid library into AWY101 cells as described above. We recovered transformants in a molten SDCAA agarose gel (1.71 g/L YNB without amino acids and ammonium sulfate (Sigma-Aldrich #Y1251), 5 g/L ammonium sulfate (Sigma-Aldrich, St. Louis, MO, #A4418), 2% dextrose (VWR #90000–904), 5 g/L Bacto casamino acids (VWR #223050), 100 g/L ampicillin (VWR # V0339)) containing 0.35% SeaPrep agarose (VWR #12001–922) spread into a thin layer (about 1 cm deep). The mixture was placed at 4°C for three hours, after which it was incubated at 30°C for 48 hours. From five aliquots, we obtained ∼1.2 million colonies. After mixing the flasks by shaking for 1 hr, we grew cells in 5 mL tubes of liquid SDCAA for five generations and stored the saturated culture in 1 mL aliquots supplemented with 5% glycerol at -80°C.

### High-throughput binding affinity assay (Tite-Seq)

Tite-Seq was performed as previously described ^12,14,15^. We performed three replicates of the assay on different days. In the first two replicates, a small portion of the library variants contained an off-target mutation (E484W) instead of the intended mutation (E484A). These variants were removed from the data analysis, and in the third replicate the library was supplemented with variants containing the intended mutation (E484A).

#### Preparation

First, we thawed yeast RBD libraries, as well as Wuhan Hu-1 and Omicron BA.1 clonal strains, by inoculating 150 μL of corresponding glycerol stock (saturated culture with 5% glycerol stored at -80°C) in 5 mL SDCAA at 30°C for 20 hr. On the next day, yeast cultures were diluted to OD600=0.67 in 5 mL SGDCAA (6.7 g/L YNB without amino acid VWR #90004-150), 5 g/L ammonium sulfate (Sigma-Aldrich #A4418), 2% galactose (Sigma-Aldrich #G0625), 0.1% dextrose (VWR #90000–904), 5 g/L Bacto casamino acids (VWR #223050), 1.065 g/L MES buffer (Cayman Chemical, Ann Arbor, MI, #70310), 100 g/L ampicillin (VWR # V0339)), and rotated at room temperature for 16–20 hr.

#### Labeling

After overnight induction, yeast cultures were pelleted, washed twice with 0.01% PBSA (VWR #45001–130; GoldBio, St. Louis, MO, #A-420–50), and resuspended to an OD600 of 1. A total of 500-700 μL of OD1 yeast cells were labeled with biotinylated human ACE2 (Acrobiosystems #AC2-H2H82E6) at each of the twelve ACE2 concentrations (half-log increments spanning 10 ^-12.5^ – 10 ^-7^ M), with volumes adjusted to limit ligand depletion effects to be less than 10% (assuming 50,000 surface RBD/cell ^33^). Yeast-ACE2 mixtures were incubated and rotated at room temperature for 20 hr. Following the incubation, yeast-ACE2 complexes were pelleted by spinning at 3000 x g for 10 minutes at 4°C, washed twice with 0.5% PBSA + 2mM EDTA, and subsequently labeled with Streptavidin-RPE (1:100, Thermo Fisher #S866) and anti-cMyc-FITC (1:50, Miltenyi Biotec, Somerville, MA, #130-116-485) at 4°C for 45 min. After this secondary labeling, yeast were washed twice with 0.5% PBSA + 2mM EDTA and left on ice in the dark until sorting.

#### Sorting and recovery

We sorted the yeast library complex on a BD FACS Aria Illu, equipped with 405 nm, 440 nm, 488 nm, 561 nm, and 635 nm lasers, and an 85 micron fixed nozzle. To minimize the spectral overlap effects, we determined compensation between FITC and PE using single-fluorophore controls. Single cells were first gated by FSC vs SSC and then sorted by either expression (FITC) or binding (PE) fluorescence. At least one million cells were sorted for each sample. In the expression sorts, singlets (based on FSC vs SSC) were sorted into eight equivalent log-spaced FITC bins. For the binding sorts, FITC+ cells were sorted into 4 PE bins (the PE-population comprised bin 1, and the PE+ population was split into three equivalent log-spaced bins 2–4 as described in Phillips and Lawrence et al. 2021). Sorted cells were collected in polypropylene tubes coated and filled with 1 mL YPD supplemented with 1% BSA. Upon recovery, cells were pelleted by spinning at 3000 x g for 10 min and resuspended in 4 mL SDCAA. The cultures were rotated at 30°C until late-log phase (OD600 = 0.9-1.4).

#### Sequencing library preparation

1.5 mL of late-log yeast cultures was pelleted and stored at -20C for at least six hours prior to extraction. Yeast display plasmids were extracted using Zymo Yeast Plasmid Miniprep II (Zymo Research # D2004), following the manufacturer’s instructions, and eluted in a 17 μL elution buffer. RBD amplicon sequencing libraries were prepared by a two-step PCR as previously described ^15,34^. In the first PCR, unique molecular identifiers (UMI), inline indices, and partial Illumina adapters were appended to the sequence library through 7 amplification cycles to minimize PCR amplification bias. We used 5 μL plasmid DNA as template in a 25 μL reaction volume with Q5 polymerase according to the manufacturer’s protocol (NEB # M0491L). Reaction was incubated in a thermocycler with the following program: 1. 60 s at 98°C, 2. 10 s at 98°C, 3. 30 s at 66°C, 4. 30 s at 72°C, 5. GOTO 2, 6x, 6. 60 s at 72°C. Shortly after the reaction completed, we added 25 μL water into reactions and performed a 1.2X magnetic bead cleanup (Aline Biosciences #C-1003–5). The purified products were then eluted in 35 μL elution buffer. In the second PCR, the remainder of the Illumina adapter and sample-specific Illumina i5 and i7 indices were appended through 35 amplification cycles (see https://github.com/desai-lab/compen-satory_epistasis_omicron/tree/main/Supplementary_Files for primer sequences). We used 33 μL of the purified PCR1 product as template, in a total volume of 50 μL using Kapa polymerase (Kapa Biosystems #KK2502) according to the manufacturer’s instructions. We incubated this second reaction in a thermocycler with the following program: 1. 30 s at 98°C, 2. 20 s at 98°C, 3. 30 s at 62°C, 4. 30 s at 72°C, 5. GOTO 2, 34x, 6. 300 s at 72°C. The resulting sequencing libraries were purified using 0.85X Aline beads, amplicon size was verified to be ∼500 bp by running on a 1% agarose gel, and amplicon concentration was quantified by fluorescent DNA-binding dye (Biotium, Fremont, CA, #31068, per manufacturer’s instructions) on Spectramax i3. We then pooled the amplicon libraries according to the number of cells sorted and further size-selected this pool by a two-sided Aline bead purification (0.5–0.9X). The final pool size was verified by Tapestation 5000 HS and 1000 HS. Final sequencing library was quantitated by Qubit fluorometer and sequenced on an Illumina NovaSeq SP with 10% PhiX.

### Sequence data processing

We processed our raw demultiplexed sequencing reads to identify and extract the indexes and mutational sites. To do so, we developed a snakemake pipeline (https://github.com/desai-lab/compensatory_epistasis_omicron/tree/main/Titeseq) that first parsed through all fastq files and separated the reads according to inline indices, UMIs, and sequence reads using Python library regex ^35^. We accepted sequences that match the entire read (with no restrictions on bases at mutational sites) within 10% bp mismatch tolerance. Next, we discarded incorrect inline indices (according to the corresponding i5/i7 indices) and parsed read sequences into binary genotypes (‘0’ for Wuhan Hu-1 allele or ‘1’ for Omicon BA.1 allele at each mutation position). Reads with errors at mutation sites (i.e. not matching either Wuhan Hu-1 allele or Omicron BA.1 allele) were discarded. Finally, we counted the number of distinct UMIs for each genotype, and collated genotype counts from all samples into a single table. The mean coverage across all replicates was ∼150x.

To fit the binding dissociation constants *K*_D,app_ for each genotype, we followed the same procedure as previously described ^21,22^. In brief, we used sequencing and flow cytometry data to calculate the mean log-fluorescence of each genotype *s* at each concentration *c*, following:

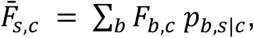

where *F* _*b,c*_is the mean log-fluorescence of bin *b* at concentration *c*, and *p* _*b,s|c*_ is the inferred proportion of cells from genotype *s* that are sorted into bin *b* at concentration *c*. The *p* _*b,s|c*_ is in turn estimated from the read counts as

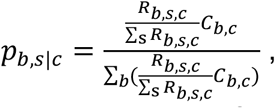

where *R* _*b, s,c*_ is the number of reads from genotype *s* that are found in bin *b* at concentration *c*, whereas *C* _*b,c*_ refers to the number of cells sorted into bin *b* at concentration *c*.

To propagate the uncertainty in the mean bin estimate, we used the formula

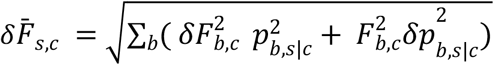

where *δ F* _*b,c*_ is the spread of log fluorescence of cells sorted into bin *b* at concentration *c*. As previously investigated, we found that estimating *δF* _*b,c*_ ≈ *σF* _*b,c*_ is sufficient to capture the variation we observed in log-fluorescence within each bin. In contrast, the error in *p* _*b*,s|*c*_ emerges from the sampling error, which can be approximated as a Poisson process when read counts are high enough.

Thus we have:

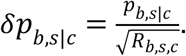

Finally, we inferred the binding dissociation constant (*K*_D,s_) for each variant by fitting the logarithm of Hill function to the mean log-fluorescence 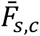, as a function of ACE2 concentrations *c*:

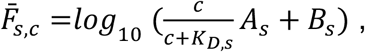

where *A* _s_ is the increase in fluorescence at ACE2 saturation, and *B* _s_ is the background fluorescence level. The fit was performed using the *curve_fit* function in the Python package *scipy*.*optimize*. Across all genotypes, we gave reasonable bounds on the values of *A* _s_ to be 10^2^-10^6^, *B* _*s*_ to be 1-10^5^, and *K*_D,s_ to be 10^−14^-10^−5^. We then averaged the inferred *K*_D,s_ values across the three replicates after removing values with poor fit (*r* ^’^ < 0.8).

### Isogenic measurements for validation

We validated our high-throughput binding affinity method by selecting 10 specific RBD clones for lower-throughput validation: Wuhan Hu-1, Omicron, 5 single-mutants (K417N, S477N, T478K, Q498R, N501), two double mutants (Q498R/N501Y and E484A/Q498R), and one genotype with four mutations (K417N/E484A/Q498R/N501Y). For each isogenic titration curve, we followed the same labeling strategy, titrating ACE2 at concentrations ranging from 10^− 12^-10^−7^ M for isogenic yeast strains that display only the sequence of interest. The mean log fluorescence was measured using a BD LSR Fortessa cell analyzer. We directly computed the mean and variances of these distributions for each concentration and used them to infer the value of - log_10_(*K*_D_) using formula (shown above) (see Extended Data Figure 1).

### Epistasis analysis

We first used a simple linear model where the effects of combinations of mutations sum to the phenotype of a sequence. The logarithm of the binding affinity log_10_(*K*_D,s_) is proportional to free energy changes, hence in a model without interaction, they would combine additively ^35^. The full K-order model can be written:

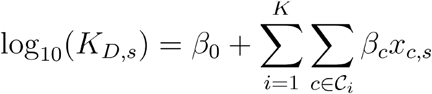

where C_i_ contains all 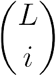 combinations of size i of the mutations and *x*_*c,s*_ is equal to 1 if the sequence contains all the mutations in and to 0 otherwise. This choice is called ‘biochemical’ or ‘local’ epistasis ^36^ and is the one used in the main text. Another option, called ‘statistical’ or ‘ensemble’ consists in replacing the coefficients *x*_*s*_ by *y*_*s*_ = 2*x*_*s*_ − 1 ∈ {−1, 1}. We present the result of this analysis, and the differences with the biochemical model, in Extended Data Figure 5.

To choose the optimal value of K, we follow the method detailed in Phillips and Lawrence et al., 2021 ^36^. Briefly, we use 10-fold cross-validation to test all values of K ≤ 6. For each value of K, the data is split into ten and each of the ten sub-dataset is used as a test set for a model trained on the rest of the data. We chose the value of K that maximizes the prediction performance (R^2^) averaged over all ten testing datasets. For this dataset we found an optimal value of K=5 (Extended Data Figure 4). Finally, we trained a K=5 model over the complete dataset to get the final coefficients. The number of parameters of the final model (~5000) is much lower than the number of observed data points (2^15^ = 32768).

As mentioned above, the logarithm of binding affinity is proportional to a free energy change, an extensive quantity. This theoretically justifies the use of a linear model. Nonetheless, in some scenarios, the interactions between mutations can be better explained by a nonlinear function with few parameters acting on the full phenotype (“global epistasis”) rather than a large number of small-effects interactions at high order (“idiosyncratic epistasis”). Our implementation is similar to that described by Sailer and Harms, 2017 ^37^ and follows closely Phillips and Lawrence et al., 2021 ^15,24^. In short, we use a logistic function Φ, with four parameters, to fit the expression:

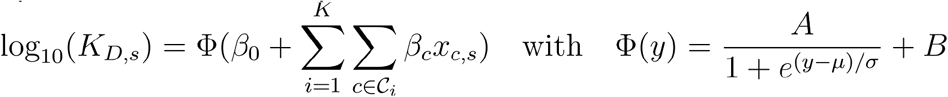

The choice of a logistic function is justified by the general form of *K*_D,app_ distribution, which slightly “plateaued” at strong *K*_D,app_. This effect is not caused by experimental artifacts (Extended Data Figure 2) but instead by a form of “diminishing returns” epistasis ^37^. Practically, the parameters are inferred by fitting successively the additive βi and the nonlinear function parameters. Although the global epistasis transformation does improve the fit, the additive coefficients observed at low order do not change significantly (Extended Data Figure 6).

### Structural analysis

We used the reference structure of a 2.79 Å cryo-EM structure of Omicron BA.1 complexed with ACE2 (PDB ID: 7WPB). In Figure 2C, the contact surface area is determined by using ChimeraX ^38^ to measure the buried surface area between ACE2 and each mutated residue in the RBD (*measure buriedarea* function, default probeRadius of 1.4Å). In Figure 2E, the distance between α-carbons is measured using PyMol ^39^.

### Order of mutations

ACE2 binding affinity impacts the fitness of SARS-CoV-2 variants and can thus be leveraged to partially infer its past trajectory. This piece of information is particularly important for Omicron BA.1, where phylogenetic information is limited. Because our dataset contains the ACE2 affinity of all possible evolutionary intermediates, we can infer the likelihoods of all pathways between the ancestral Wuhan Hu-1 sequence and Omicron BA.1. To do this we need to choose a selection model. The circumstances in which the Omicron variant evolved are unknown, and the evolutionary fitness of the virus is more complex than its capacity to bind ACE2 – immune pressure, structural stability, and expression level also play a role, among many other factors ^40^. In addition, back-mutations are common in viral evolution and selection pressure can change depending on whether the strain is switching hosts rapidly or part of a long-term infection. Here, we have chosen to adopt an extremely simple weak-mutation/strong-selection regime of viral evolution.

In that model, selection proceeds as a Markov process, where the population is characterized by a single sequence that acquires a single mutation at each discrete step ^29,41^. We assume that back mutations (i.e. a residue changing from the Wuhan Hu-1 amino-acid to the BA.1 one) are not possible. Once such a sequence is generated, it will either fix in the full population or die out. The important parameter is then the fixation probability, which depends on the binding affinity of both the original and mutated sequences.

We choose to use the commonly used classical fixation probability ^42^, for a mutation with selection coefficient σ in a population of size N:

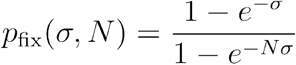

Here, the selection coefficient is proportional to the difference in log binding affinities between the two sequences. We use this model in the “strong selection” limit (N → ∞ and σ →∞), where a mutation fixes if it is advantageous or if it is the less deleterious choice among all the leftover mutations. Weaker selection models give qualitatively similar results. In terms of implementation, we use a transition matrix approach that allows us to quickly compute the probability that each residue appears at a specific position.

### Force directed layout

The high-dimensional binding affinity landscape can be projected in two dimensions with a force-directed graph layout approach (see https://desai-lab.github.io/wuhan_to_omicron/). Each sequence in the antibody library is a node, connected by edges to its single-mutation neighbors. An edge between two sequences s and t is given the weight:

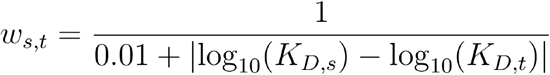

In a force-directed representation, nodes repel each other, while the edges pull together the nodes they are attached to. In our scenario, this means that nodes with a similar genotype (a few mutations apart) and a similar phenotype (binding affinity) will be close to each other in two dimensions.

Importantly this is not a “landscape” representation: the distance between two points is unrelated to how easy it is to reach one genotype from another in a particular selection model. Practically, after assigning all edge weights, we use the layout function *layout_drl* from the Python package *iGraph*, with default settings, to obtain the layout coordinates for each variant.

### Genomic data

To analyze SARS-CoV-2 phylogeny (Figure 4A and 4B), we used all complete RBD sequences from all SARS-CoV-2 genomes deposited in the Global Initiative on Sharing All Influenza Data (GISAID) repository ^43–45^ with the GISAID Audacity global phylogeny (EPI_SET ID: EPI_SET_20220615uq, available on GISAID up to June 15, 2022, and accessible at https://doi.org/10.55876/gis8.220615uq). We pruned the tree to remove all sequences with RBD not matching any of the possible intermediates between Wuhan Hu-1 and Omicron BA.1 and analyzed this tree using the python toolkit ete3 ^46^. We measured the frequency of each mutation (Figure 4A) by counting how many times it occurs independently in the tree (i.e., how often the mutation appears on a node whose parent node does not have that mutation). For Figure 4B, we counted two mutations as co-appearing if both mutations are absent in the parent node and contained in at least one of the descendant nodes. This strategy of studying the relative frequency of co-appearing mutations is a specific case to the method developed in Kryazhimskiy et al ^41^, which infers epistasis between mutations from phylogenetic data (the general method was not applicable in this specific dataset due to its size).

### Statistical analyses and visualization

All data processing and statistical analyses were performed using R v4.1.0 ^47^ and python 3.10.0 ^48^. All figures were generated using ggplot2 ^49^ and matplotlib ^50^.

## ACKNOWLEDGEMENTS

We thank Zach Niziolek for assistance with flow cytometry and members of the Desai lab for helpful discussions. T.D. acknowledges support from the Human Frontier Science Program Postdoctoral Fellowship, A.M.P. acknowledges support from the Howard Hughes Medical Institute Hanna H. Gray Postdoctoral Fellowship, J.C. acknowledges support from the National Science Foundation Graduate Research Fellowship, and M.M.D. acknowledges support from the NSF-Simons Center for Mathematical and Statistical Analysis of Biology at Harvard University, supported by NSF grant no. DMS-1764269, and the Harvard FAS Quantitative Biology Initiative, grant DEB-1655960 from the NSF and grant GM104239 from the NIH. J.D.B. acknowledges support from NIH/NIAID grant R01AI141707 and is an Investigator of the Howard Hughes Medical Institute. We gratefully acknowledge all data contributors, i.e. the Authors and their Originating laboratories responsible for obtaining the specimens, and their Submitting laboratories for generating the genetic sequence and metadata and sharing via the GISAID Initiative. Computational work was performed on the FASRC Cannon cluster supported by the FAS Division of Science Research Computing Group at Harvard University.

## AUTHOR CONTRIBUTIONS

Conceptualization: A.M., T.D., A.M.P., J.C., T.N.S., A.J.G., J.D.B., and M.M.D. Methodology: A.M., T.D., A.M.P., J.C., S.N., T.N.S., and A.J.G. Library design and production: A.M., T.D., A.M.P., J.C., and A.J.G. Experiments: A.M., T.D., A.M.P., J.C., and A.A.R. Validation: A.M., T.D., A.M.P., J.C., S.N., and T.N.S. Data analysis: A.M., T.D., A.M.P., J.C., S.N., and T.N.S. Supervision: A.M.P, J.D.B., and M.M.D. Funding acquisition: J.D.B. and M.M.D. Writing—original draft: A.M., T.D., A.M.P., J.C., and M.M.D. All the authors reviewed and edited the manuscript.

## MATERIALS AND CORRESPONDENCE

Correspondence and requests for materials should be addressed to M.M.D.

## DATA AND CODE AVAILABILITY STATEMENT

Raw sequencing reads have been deposited in the NCBI BioProject database under accession number PRJNA849979. All associated metadata are available at https://github.com/desai-lab/compensatory_epistasis_omicron.

## EXTENDED DATA FIGURES AND CAPTIONS

**Extended Data Figure 1:**
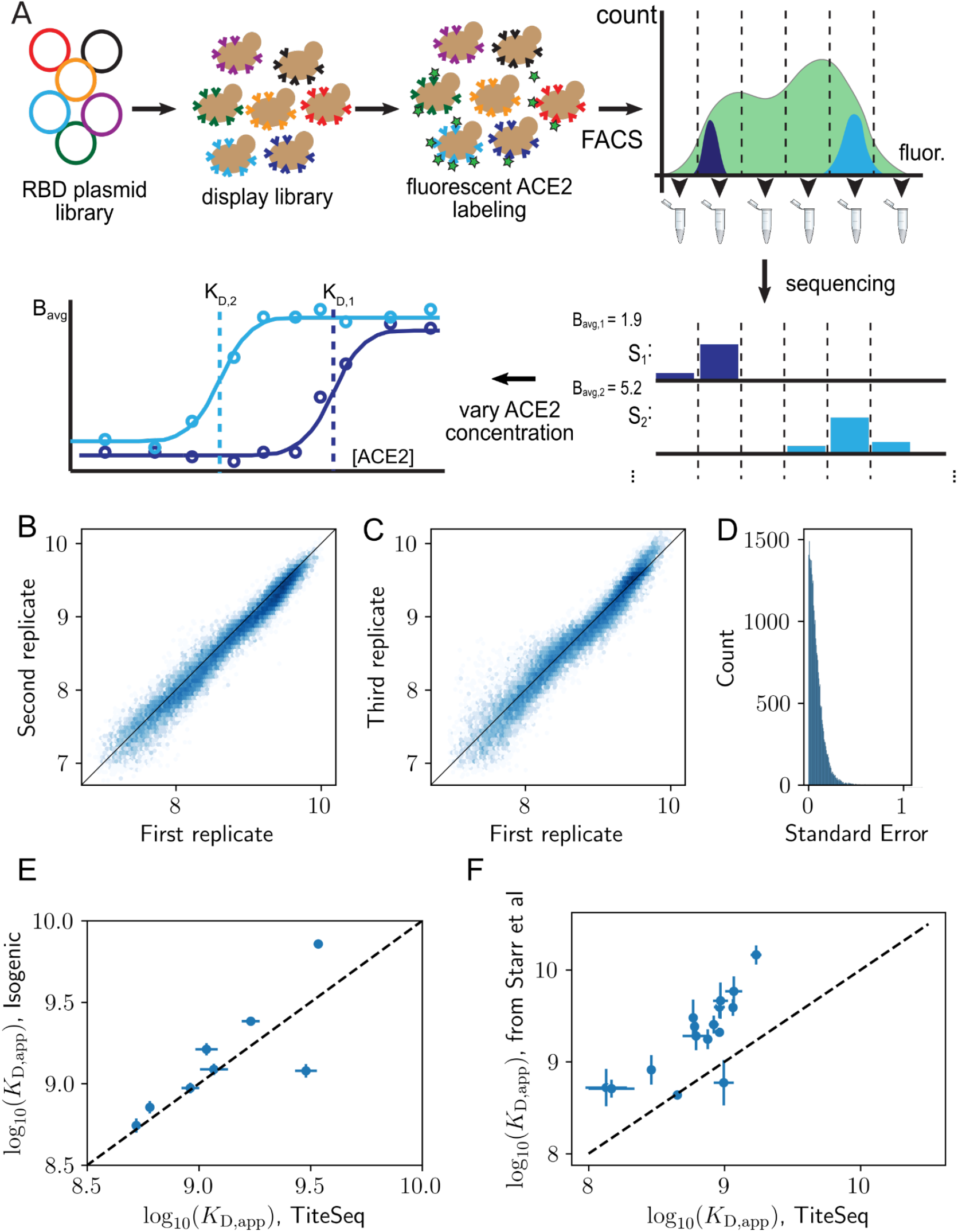
Schematic overview of the experimental method and reproducibility of dissociation constants determined via Tite-seq. **(A)** The plasmid library of RBD variants is first transformed into a standard yeast display strain. The library is incubated with soluble, fluorescent ACE2 and sorted by flow cytometry into bins based on ACE2 fluorescence. Deep sequencing of each bin yields an estimate for the mean bin (B_avg_) of each RBD variant. This is repeated for varying ACE2 concentration to produce a titration curve. Since the fluorescence is linearly related to the RBD occupancy on the yeast cell surface, apparent equilibrium dissociation constants can be inferred by fitting B_avg_ to the ACE2 concentration. (**B**) Correlation of -log(*K*_D,app_) between the first and second biological replicates. (**C**) Correlation of -log(*K*_D,app_) between the first and third biological replicates. (**D**) Distribution of the standard error of -log(*K*_D,app_) between biological replicates. **(E)** Isogenic measurements (see Methods) versus Tite-Seq measurement with a 1:1 dotted line. **(F)** Comparison of Tite-Seq *K*_D_ measurements with independent *K*_D_ measurements reported in Starr et al ^12^ with a 1:1 dotted line.

**Extended Data Figure 2:**
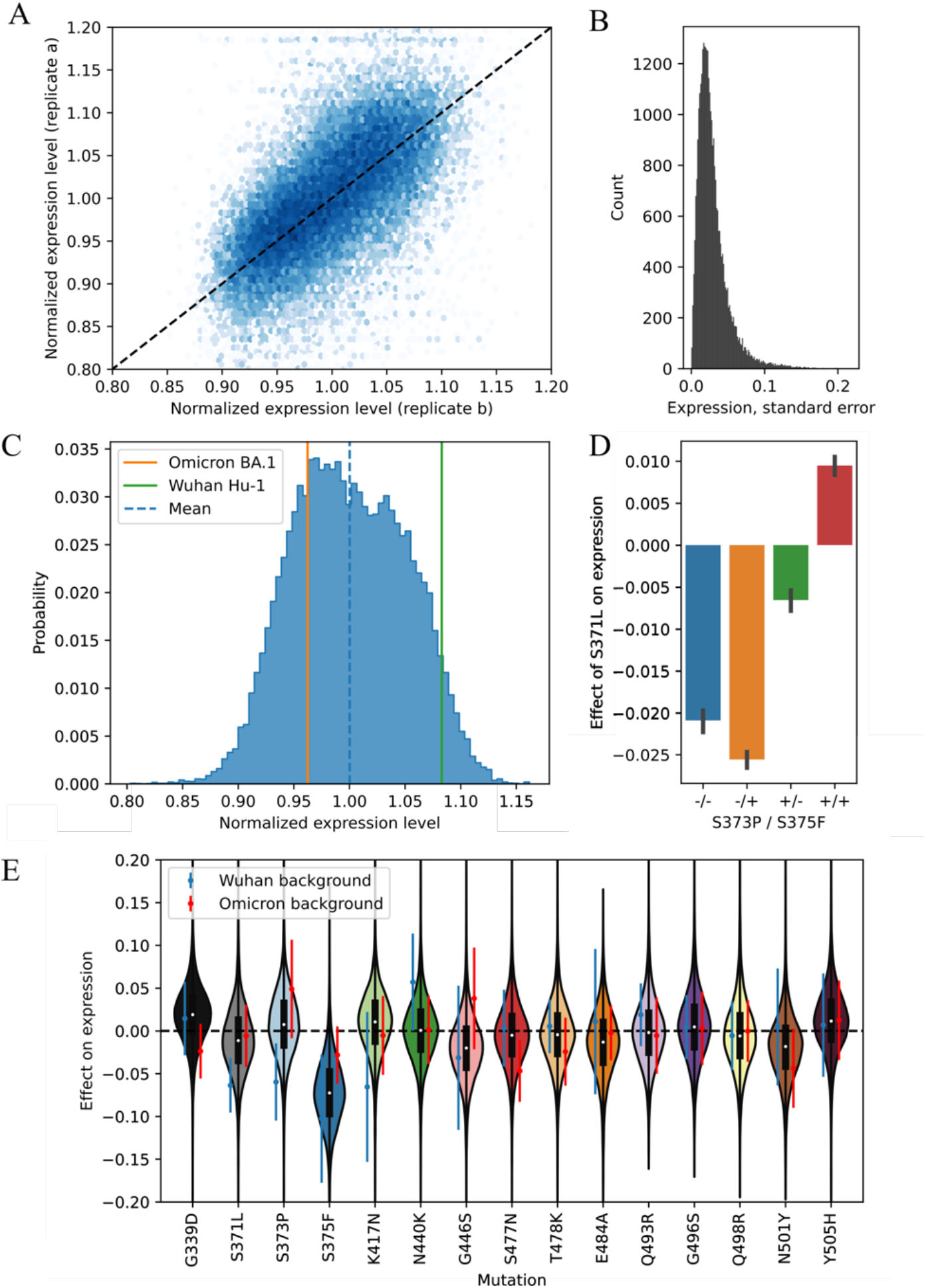
Expression level of RBD in the yeast display system. (**A**) Correlation of normalized expression levels between the first and second biological replicates. (**B**) Distribution of the normalized expression levels between biological replicates. (**C**) Distribution of the normalized yeast-display expression of each RBD variant in the library. Vertical red and green lines represent the expression for Wuhan Hu-1 and Omicron BA.1, respectively. (**D**) Effect of the S371L mutation on expression levels depending on the S373P and S375F background. **(E)** Mutational effects (defined as the difference in normalized expression after adding one mutation) for each Omicron BA.1 RBD mutation. Violin plots show full distribution of effects, where black lines indicate 25th and 75th percentiles and the black point denotes mean. Blue and red points specify effects on Wuhan Hu-1 and into Omicron BA.1 variants, respectively.

**Extended Data Figure 3:**
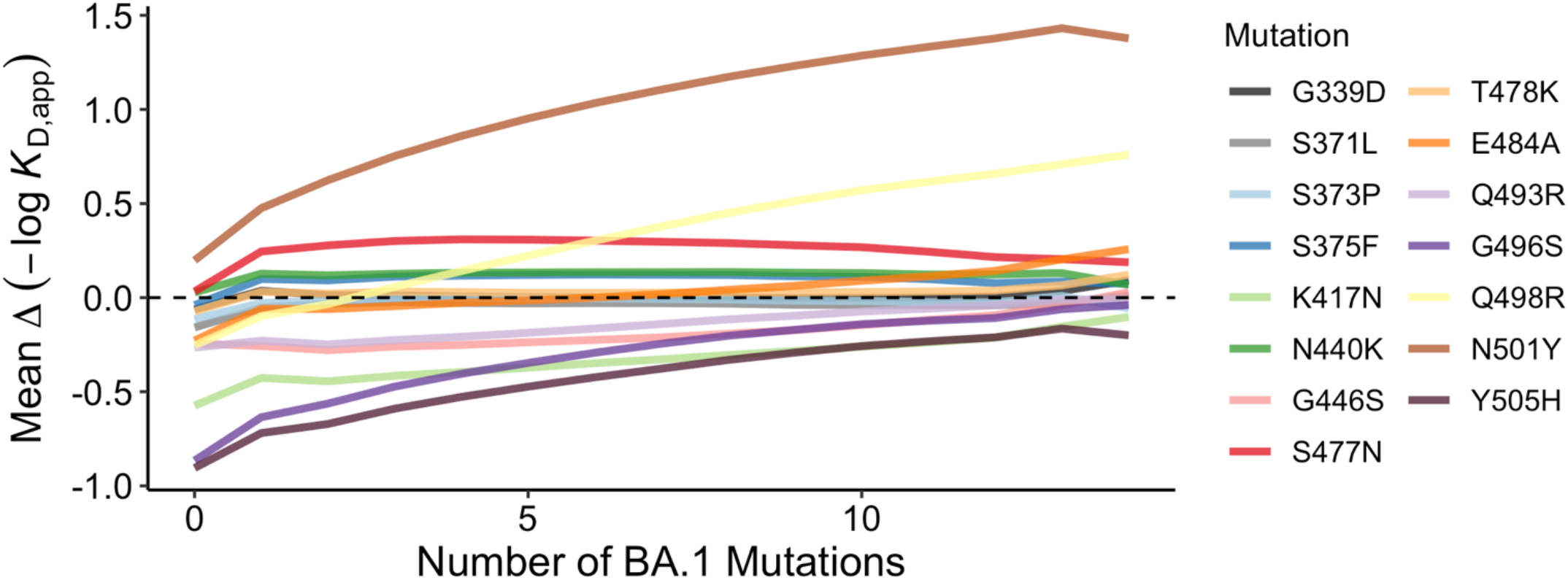
Change in ACE2 affinity across number of mutations. The mean effect of each mutation is plotted against the number of BA.1 mutations in the genotypic background. Dashed line indicates no change in affinity.

**Extended Data Figure 4:**
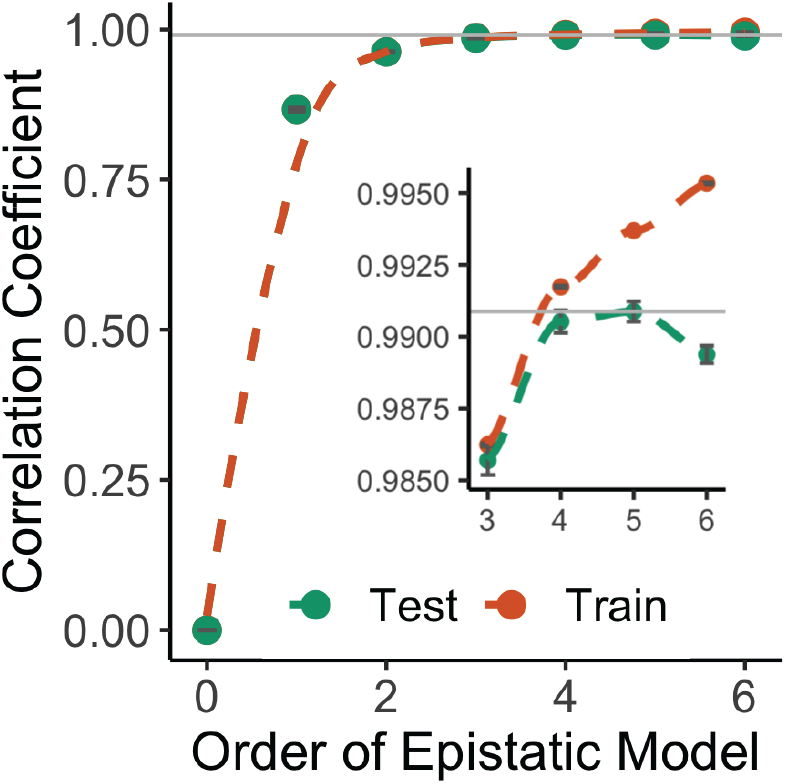
Truncation of biochemical epistasis model. Correlation coefficients between the measured values of -log(*K*_D,app_) and the model estimate for various orders of epistatic model. Correlations are computed on the subset of the dataset on which the model was trained (orange) and on the hold-out subset (green), averaged over the 10 folds of cross-validation. The inset is a zoomed-in version for orders 3 to 6.

**Extended Data Figure 5:**
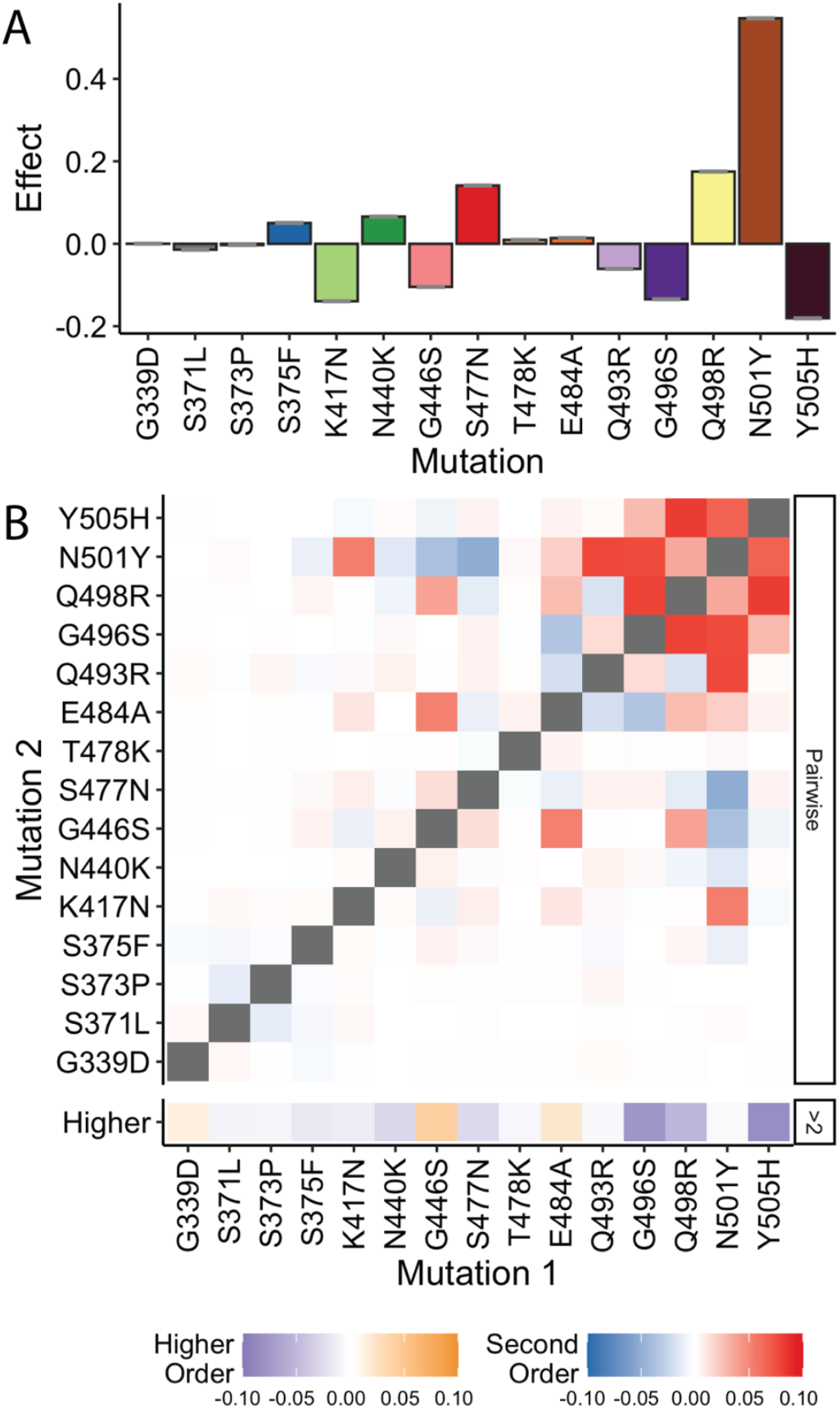
Alternative model of statistical epistasis. **(A)** Linear effect of each mutation in the statistical epistasis model that is truncated at the fourth order. **(B)** Second-order epistatic interaction coefficients and higher order interaction in the statistical epistasis model.

**Extended Data Figure 6:**
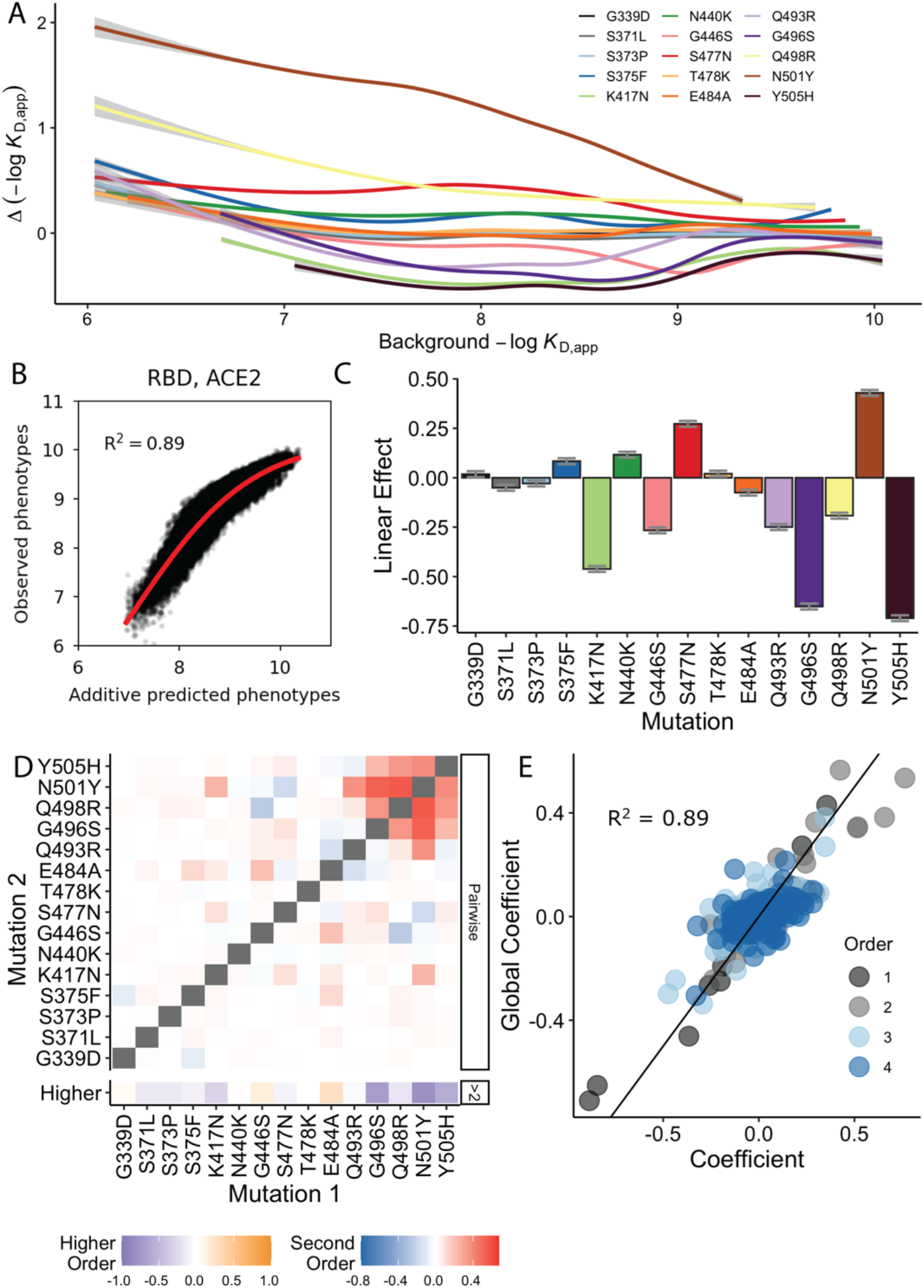
Global epistasis. (**A**) Relationship between the binding affinity and the mean effect of an additional mutation on this background. **(B)** Relationship between the observed binding affinity and the affinity predicted with a linear additive model without epistasis. The red line represents the global epistasis function. **(C)** Linear effect of each mutation in the global epistasis model that is truncated at the fourth order. (**D**) Second-order and higher-order epistatic interaction coefficients in the global epistatic model. **(E)** Correlation between the epistatic interaction coefficients of the models with and without global epistasis. The black line represents the best fit.

**Extended Data Figure 7:**
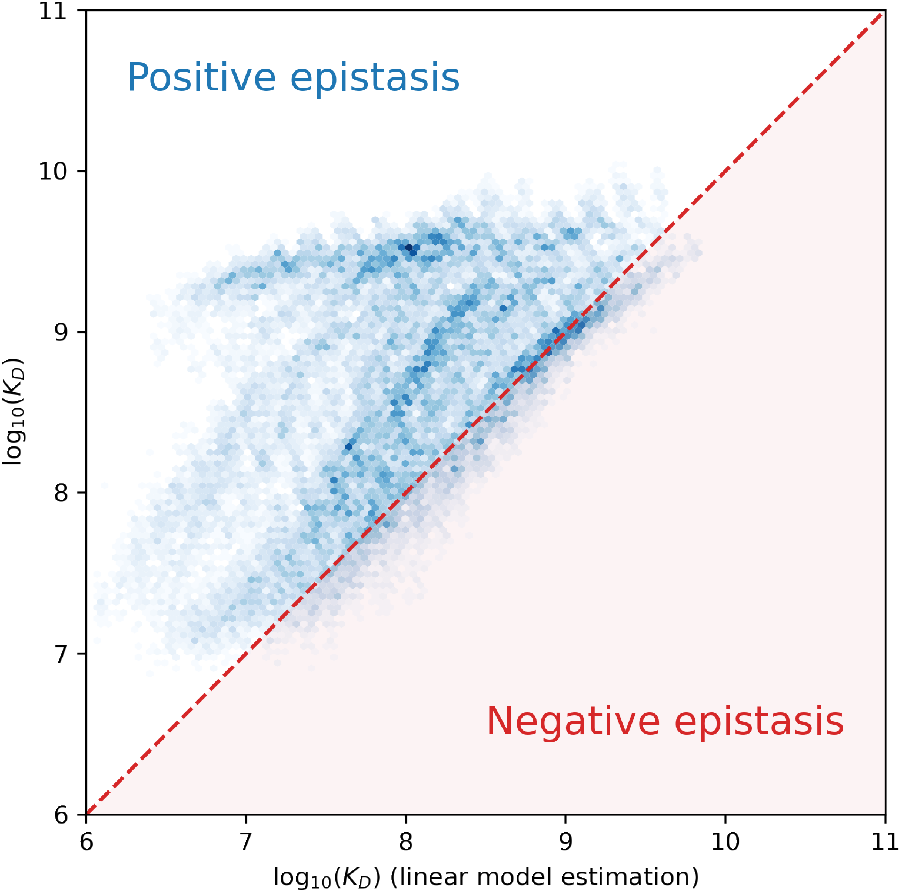
Comparison between the linear model estimate of the binding affinity and the measured binding affinity. The x-axis is the predicted binding affinity, using only the linear coefficients of the full 5th-order model; the y-axis is the measured binding affinity.

**Extended Data Figure 8:**
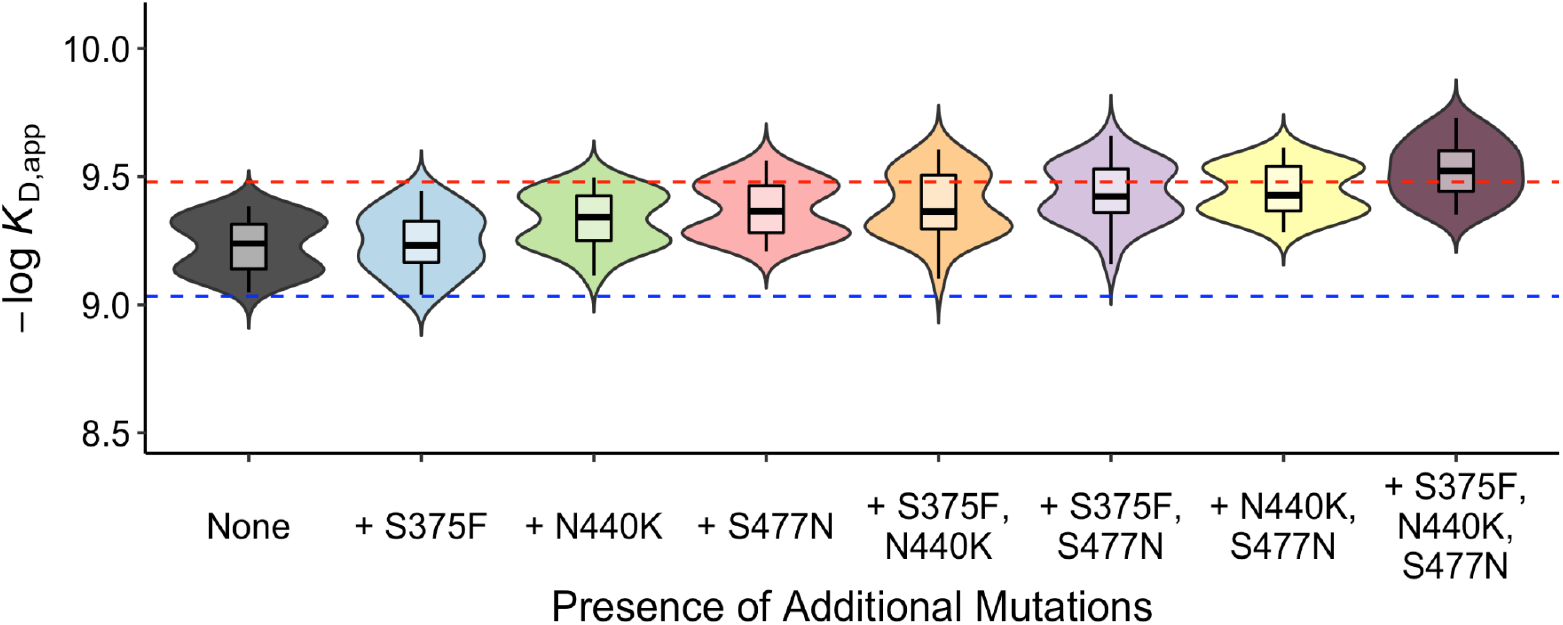
Binding affinity of escape genotypes with additional compensatory mutations. The ACE2 binding affinities of variants with all seven mutations discussed in the main text (the five escape mutations K417N, G446S, E484A, Q493R, and G496S, plus Q498R and N501Y) with all possible combinations of three other mutations (S375F, N440K, and S477N). Blue and red dashed lines represent Wuhan Hu-1 and Omicron BA.1 affinity, respectively.

**Extended Data Figure 9:**
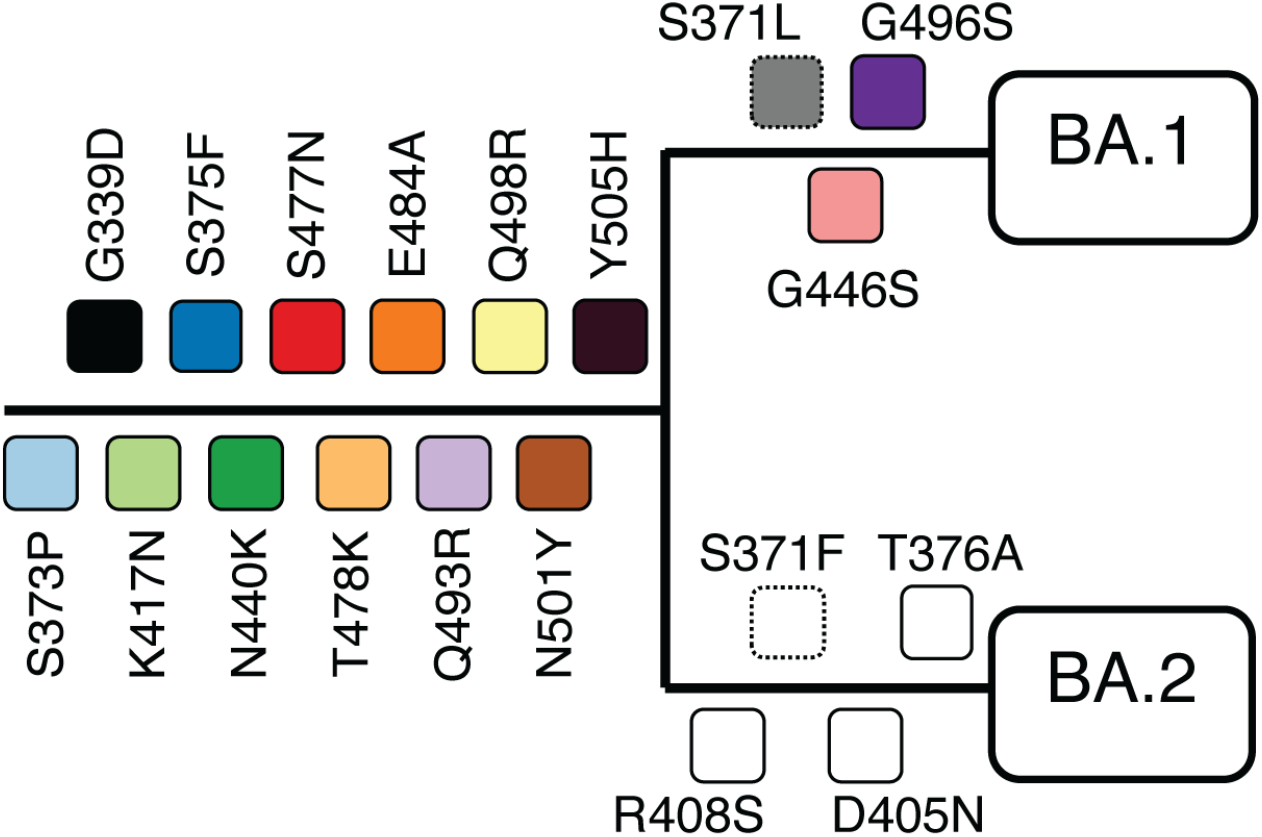
Phylogeny of BA.1 and BA.2 showing mutations in spike protein RBD. Mutations are colored as in Figure 2A. Dashed boxes indicate mutations with ambiguous positions on the tree.

**Extended Data Figure 10:**
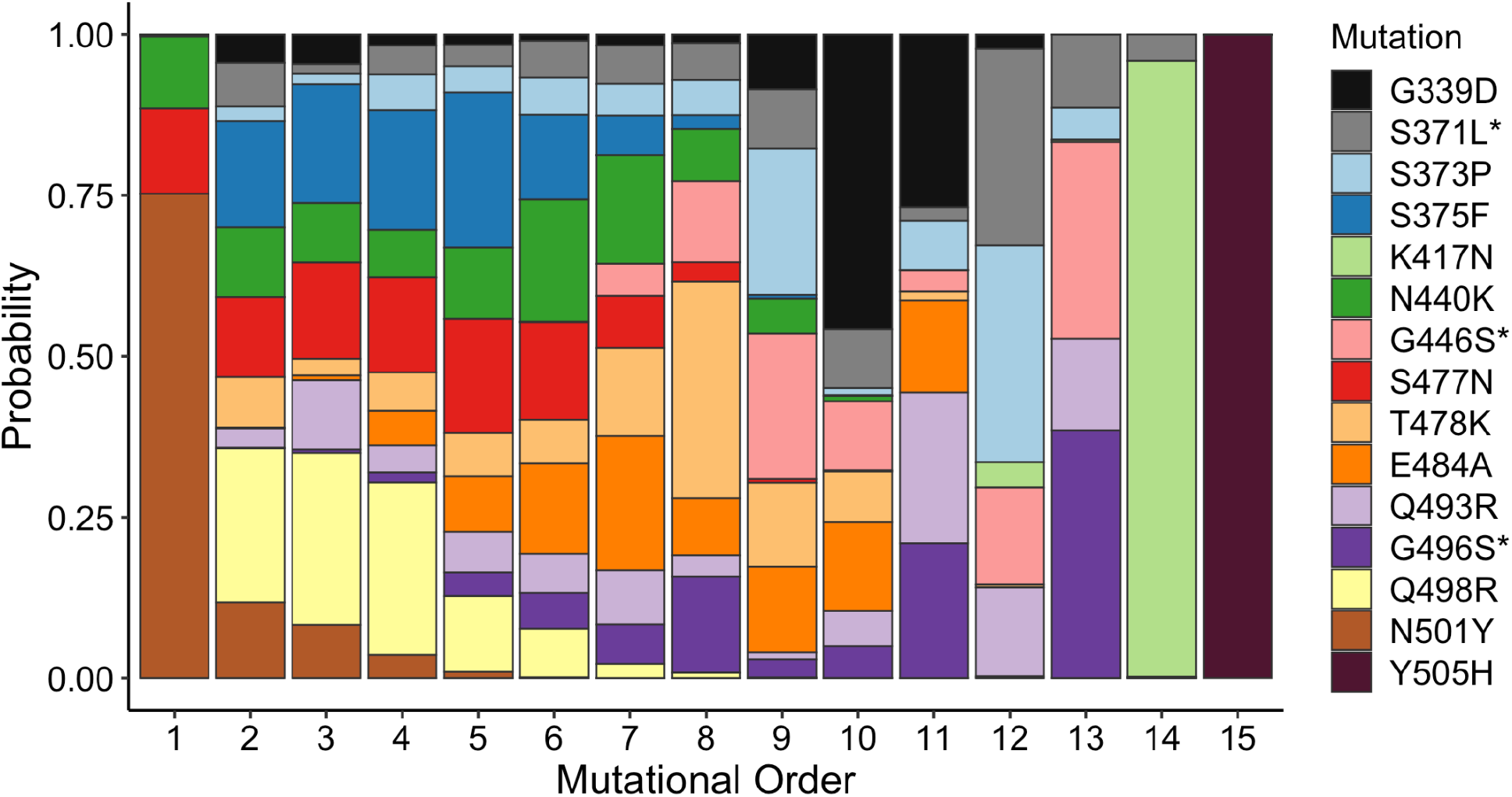
Inferred order of mutations. Conditional probability of mutation order from Wuhan-Hu-1 to Omicron BA.1 variant, assuming a classical population dynamics model (see Methods). Mutations with asterisks are known to happen last (see Extended Data Figure 9).

